# SHP-1 interacts with NFκB1 to inhibit its phosphorylation and nuclear translocation to suppress excessive bacterial inflammation

**DOI:** 10.1101/2024.04.13.589348

**Authors:** Ningning Wang, Suxu Tan, Muyuan Wang, Hongning Liu, Sen Han, Zhendong Wu, Jie Ma, Songlin Chen, Zhenxia Sha

**Author notes:** Corresponding author. (Z. S.); (S. C.). These authors contributed equally to this work (Ningning Wang, Suxu Tan).

## Abstract

The protein tyrosine phosphatase SHP-1 is a key negative regulator in cancer by dephosphorylating multiple target molecules. Specially in the NFκB signaling, where NFκB1/Rela dimer translocate to the nucleus and activate target gene transcription, SHP-1 inhibits the phosphorylation of Rela, while its regulation on NFκB1 has been unknown, especially in pathogen-induced inflammation. Chinese tongue sole, a representative flatfish, has been widely used as a genomics and disease model. Using the teleost and cellular model, we revealed for the first time that SHP-1 inhibits NFκB1 phosphorylation and nuclear translocation by interacting with NFκB1, thereby suppressing NFκB signaling to inhibit bacterial inflammation. In addition, we showed that SHP-1 decreased mortality and alleviated histopathological deterioration, manifested in the inhibition of immune-related pathways and secretion of pro-inflammatory cytokines. Using cellular model, SHP-1 overexpression reduced macrophages M1 polarization, phagocytosis, and oxidative stress, while silencing SHP- 1 exhibited opposite effects. Our findings systematically dissect the functions of SHP- 1 and provide mechanistic insights into the control of inflammation-related diseases.

**Teaser:** SHP-1 help maintain the cellular and individual homeostasis by inhibiting the excessive inflammation and immunity via regulating the NFκB signaling.

## Introduction

SHP-1 (PTPN6) belongs to the classic non-receptor protein tyrosine phosphatase, which counterbalances the activation of protein tyrosine kinase and modulates the kinetics of signal transduction (*1*). Its expression level plays an important role in regulating protein tyrosine dephosphorylation, affecting various physiological processes, including growth, proliferation, differentiation, and disease (*2, 3*). Due to its biological significance, SHP-1 has garnered attention in the field of cancer research and shown therapeutic potential. For instance, SHP-1 dephosphorylated signal transducer and activator of transcription 3 (STAT3) to suppress the corresponding signaling in breast cancer, leading to tumor cells apoptosis (*4–6*). In lung cancer and hepatocellular carcinoma, SHP-1 acted as a tumor suppressor to inactivate phosphatidylinositol-3- kinase/protein kinase B (PI3K/AKT), STAT3 and nuclear factor kappa-B (NFκB) signaling pathways (*7–9*). In addition to cancer, studies on the role of SHP-1 in inflammatory response and disease have received increasing attention, mainly using murine model. For instance, SHP-1 knockout promoted pro-inflammatory cytokine production, aggravated oxidative stress, and increased mortality in lipopolysaccharide (LPS)-induced liver endotoxemia of mouse (*10*).

Inflammation can be evoked by several causes, and infection-induced inflammation holds a significant place (*11*). When the body is infected, the immune system responds by releasing cytokines and other inflammatory mediators to fight the invading pathogens. An effective host defense against fulminant pathogen infection depends on the orchestrated balance of robust inflammatory/immune response and the timely control of these responses to avoid tissue damage and even death. Dysregulated inflammatory response can lead to the overactivation of multiple signaling pathways, including but not limited to NFκB (*12–15*), toll like receptor 5-myeloid differentiation primary response 88 (TLR5-MYD88) (*16, 17*), and STAT3 (*18–20*) signaling, resulting in the occurrence of cytokine storms with the overexpression of interleukin-1β (IL-1β), interleukin-6 (IL-6), interferon gamma (IFN-γ), and tumor necrosis factor-α (TNF-α), threatening the host’s life (*21*). These emphasize the importance of negative regulation of the overactivated inflammatory/immune response, highlighting the necessity to elaborate the role and molecular mechanisms of SHP-1 in these signaling pathways.

As lower vertebrates, fish have been used as excellent models for genetic and disease research, which can provide valuable information and clues for their usage in higher vertebrates (*22, 23*). However, in terms of SHP-1, little has been done and the underlying regulatory mechanism is unknown in fish. For instance, in gibel carp (*Carassius gibelio*), diversity immunoglobulin domain protein (DICP) genes were shown to interact with SHP-1 and SHP-2, inhibiting the inflammatory response induced by interferon (*24, 25*). In studies of channel catfish (*Ictalurus punctatus*) and rainbow trout (*Oncorhynchus mykiss*), the researchers mainly focused on the cloning and discovering inhibitory receptors on the surface of white blood cells which can recruit SHP-1 to enhance the inhibitory effect on immune responses (*26, 27*). Using Chinese tongue sole (*Cynoglossus semilaevis*), we previously demonstrated that *shp-1* was differentially expressed after the infection with *V. anguillarum* (*28*), suggesting its importance in bacterial inflammation and immune response. We therefore intend to use the same fish model and bacterium to reveal the function and mechanism of SHP-1.

In this study, we revealed the anti-inflammatory role of SHP-1 in response to bacterial infection using overexpression and silencing approaches both *in vivo* and *in vitro*, and further discovered a new mechanism underlying its function. We first demonstrated the role of SHP-1 in alleviating inflammation at the individual mortality level and pathological level. More in-depth at the molecular level, the expression of inflammatory cytokines and immune-related signaling supported and explained the phenotypic change. To validate these findings observed in individuals of Chinese tongue sole, we further constructed the LPS induced inflammation model of hepatocytes and macrophages. Using the cell platform, we showed that the manipulated SHP-1 expression affected the macrophage polarization, phagocytic activity, oxidative stress, and the expression of inflammatory cytokines and immune-related signaling. All results obtained using cells were consistent with those using individuals, showcasing the accuracy and rigor of our findings. Importantly, utilizing co-immunoprecipitation (Co-IP), immunofluorescence (IF) co-localization, and immunohistochemical (IHC) analysis, we revealed a new mechanism that SHP-1 inhibits the phosphorylation and nuclear translocation of NFκB1 via interacting with NFκB1 to regulate inflammation. These results may provide innovative approaches for therapeutic intervention in cancer and infectious inflammation.

## Results

### SHP-1 protects Chinese tongue sole against tissue damage and death after *V. anguillarum* infection

To gain an insight into the function of SHP-1 in inflammation and immune response of Chinese tongue sole, we designed the experiment by regulating the expression of SHP- 1 artificially (Fig. 1A). As described in Materials and Methods 2.1, the fish were divided into blank control (CO) group, negative control (NC) group, SHP-1 overexpression (OE) group, and SHP-1 inhibition (IN) group. Hyperemia/hemorrhage was observed in internal tissues (intestine, liver, spleen, and gill) of representative individuals in the NC and IN group, while the OE group was relatively normal (Fig. 1B). Corresponding to the tissue symptoms, the survival rate of OE group was the highest after infection with different concentrations of *V. anguillarum* (Fig. 1, C and D). After infection with *V. anguillarum* at 80% lethal dose (LD80), the mortality was 85.0% in the NC group, 61.7% in the OE group, and 86.7% in the IN group (Fig. 1C). Due to the high lethal concentration of *V. anguillarum*, the Chinese tongue sole experienced rapid death within 24 h post infection. In the experiment of *V. anguillarum* infection at median lethal dose (LD50), SHP-1 significantly reduced the mortality rate of Chinese tongue sole, with the mortality rate of NC group being 43.3%, OE group 6.7%, and IN group 40.0% (Fig. 1D). These results demonstrated that SHP-1 significantly reduced the mortality of Chinese tongue sole and alleviated the tissue damage regardless of the concentration of *V. anguillarum*.

**Fig. 1.**
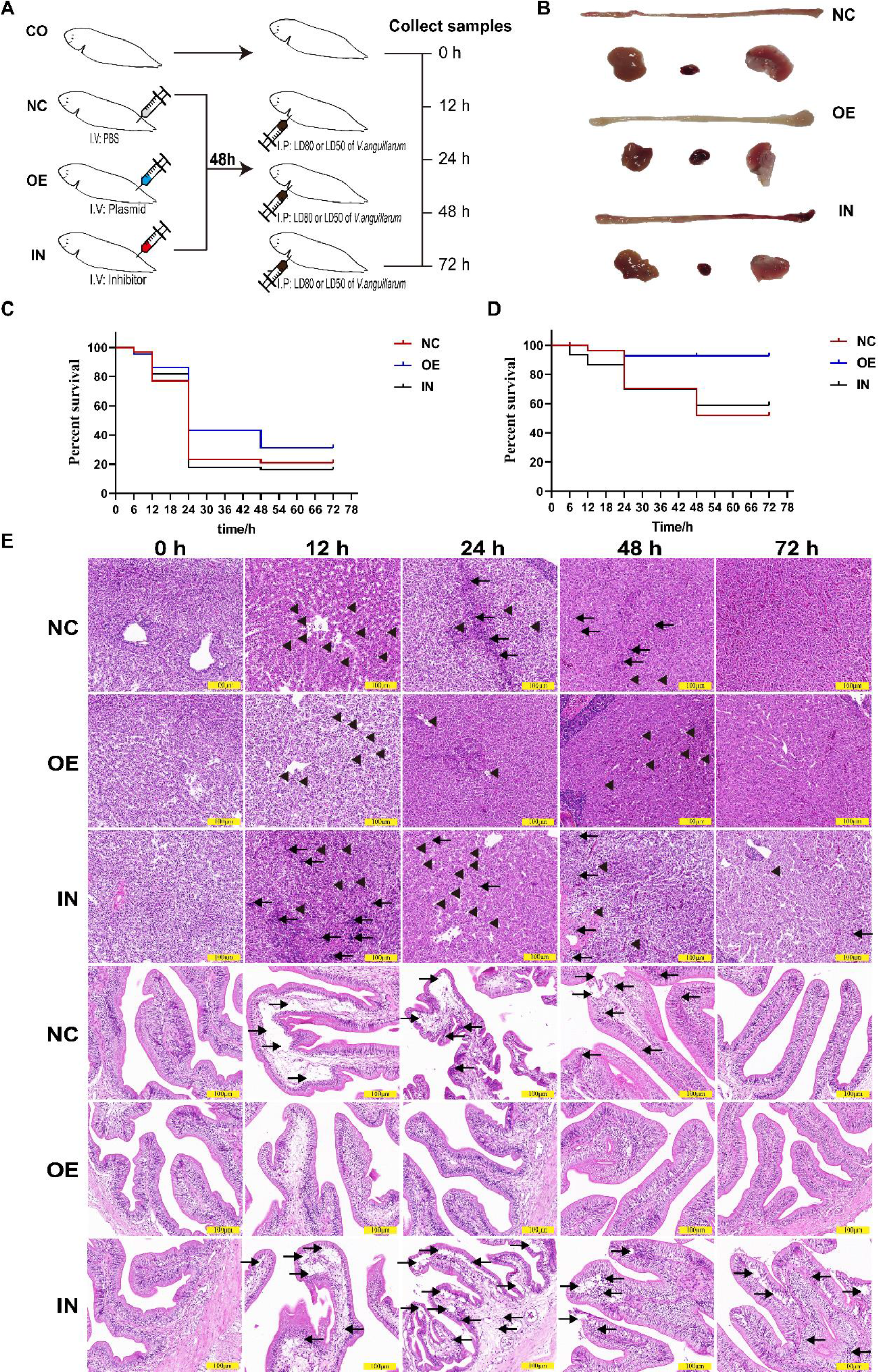
SHP-1 protects Chinese tongue sole against death and tissue damage caused by infection of *V. anguillarum*. (**A**) Schematic diagram of individual experiments on Chinese tongue sole. CO, blank control group; NC, PBS injection followed by *V. anguillarum* infection; OE, SHP-1 overexpression followed by *V. anguillarum*; IN, SHP-1 inhibition followed by *V. anguillarum* infection. (**B**) Morphological changes in the intestine, liver, spleen, and gill of Chinese tongue sole after infection with *V. anguillarum*. (**C**) Survival of Chinese tongue sole after infection with *V. anguillarum* at LD80. N = 60 individuals in each group. (**D**) Survival of Chinese tongue sole after infection with *V. anguillarum* at LD50. N = 60 individuals in each group. (**E**) Histopathological images of liver (upper three rows) and intestine (lower three rows) of Chinese tongue sole at 0 h, 12 h, 24 h, 48 h, 72 h after *V. anguillarum* infection at LD80. The long arrow pointing to the left represents inflammatory cells infiltration; The short arrow pointing to the left represents vacuolar degeneration of liver cells; The long arrow pointing to the right represents separation of intestinal mucosal layer and submucosal layer, intestinal villus rupture. Scale bars, 100 μm (E).

To reveal the role of SHP-1 at a deeper level following the observation of tissue phenotype, hematoxylin-eosin (HE) staining was performed to show the pathological damage of liver and intestine caused by *V. anguillarum* infection in different groups. As shown in Fig. 1E, inflammatory cell infiltration and vacuolar degeneration were observed in liver after infection, especially at 12-48 h post infection in the NC and IN group, while less damage was observed in the OE group, reflecting the protective effect of SHP-1. Similarly, HE staining showed severe rupture of mucosal layer and submucosal layer as well as inflammatory cell infiltration in the intestine of NC and IN group, while no significant pathological change was observed in the OE group.

### SHP-1 blocks the occurrence of cytokine storms caused by infection of *V. anguillarum*

To investigate whether SHP-1 suppresses inflammatory response by regulating inflammation related signals. In this study, qRT-PCR was conducted to measure the expression levels of *il-1β, il-6, tlr5, myd88, nfκb1, nfκb2, jak2*, and *stat3* as indicators of inflammatory response and representatives of corresponding pathways. Overall, the expression levels of these genes in each group first increased and then decreased over time (Fig. 2, A to H). At each time point, overexpression of SHP-1 generally suppressed the expression of the studied genes, while SHP-1 inhibitor increased their expression, and statistical significance was usually observed at 12 and 24 h post infection.

**Fig. 2.**
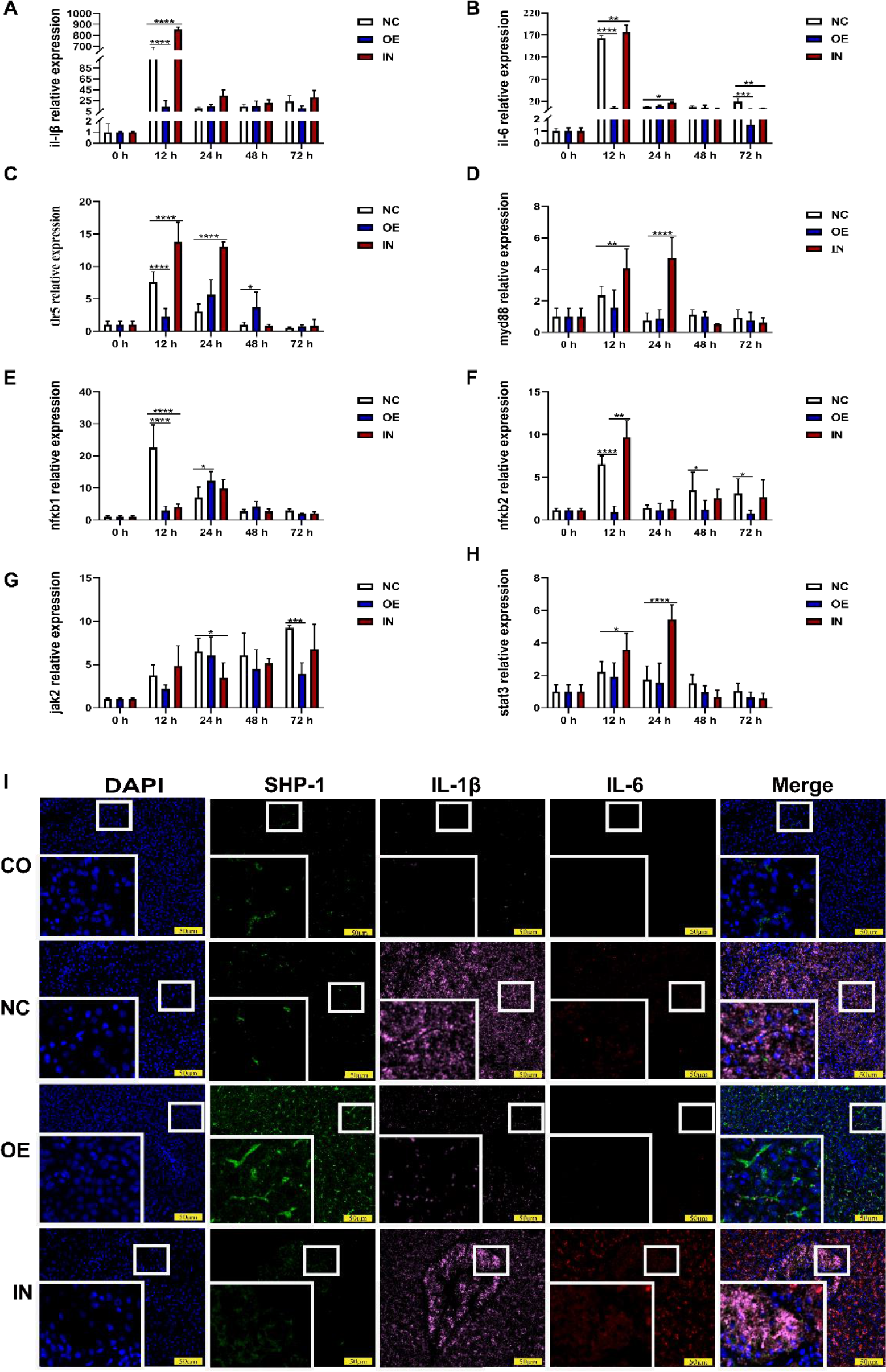
SHP-1 blocking the occurrence of cytokine storms caused by infection of *V. anguillarum*. (**A to H**) Expression of *il-1β*, *il-6, tlr5*, *myd88*, *nfκb1*, *nfκb2*, *jak2*, *stat3* in the liver of Chinese tongue sole after infection with *V. anguillarum* at LD80. (**I**) Expression of SHP-1, IL-1β, IL-6 of liver tissue by tyramide signal amplification technology. CO, blank control group; NC, PBS injection followed by *V. anguillarum* infection; OE, SHP-1 overexpression followed by *V. anguillarum*; IN, SHP-1 inhibition followed by *V. anguillarum* infection. DAPI staining was used to indicate the nuclei with blue fluorescence; SHP-1 combined with CoraLite488-conjugated goat anti-rabbit IgG secondary antibody shows green fluorescence, IL-1β combined with Cy3 goat anti- rabbit IgG secondary antibody shows pink fluorescence; IL-6 combined with Cy5 goat anti-rabbit IgG secondary antibody displays red fluorescence. Data are presented as means ± SD, n = 3 biologically independent experiments [(A) to (H)]. * *P* < 0.05, ** *P* < 0.01, *** *P* < 0.001, **** *P* < 0.0001 by one-way ANOVA. Scale bars, 50 μm (I).

IL-1β and IL-6 are important pro-inflammatory mediators, usually used as indicators of in the occurrence of cytokine storms (*29, 30*). We therefore detected the expression of SHP-1, IL-1β and IL-6 in the liver of Chinese tongue sole by immunofluorescence staining. The results exhibited excessive expression of IL-1β and IL-6 in NC and IN group, whereas SHP-1 overexpression attenuated their expression. Notably, SHP-1 expression and IL-6/IL-1β production showed opposite trends, demonstrating the role of SHP-1 in controlling inflammation (Fig. 2I).

### LPS induces inflammation in hepatocytes/macrophages and affects macrophage polarization and phagocytic activity

We further constructed LPS-induced cellular inflammation models. After LPS stimulation, the expression of SHP-1 was upregulated in both hepatocytes and macrophages by qRT-PCR and Western blot (Fig. 3, A to C, E to G). qRT-PCR results showed that 5 μg/mL LPS induced inflammation with the significantly upregulated pro-inflammatory cytokine *il-1β* and *il-6* in hepatocytes and macrophages. In addition, genes of *tlr5*, *myd88*, *nfκb1*, *nfκb2*, *jak2*, and *stat3* were also up-regulated, corresponding to the activation of TLR5-MYD88-NFκB and JAK-STAT3 signaling pathway (Fig. 3, D and H).

**Fig. 3.**
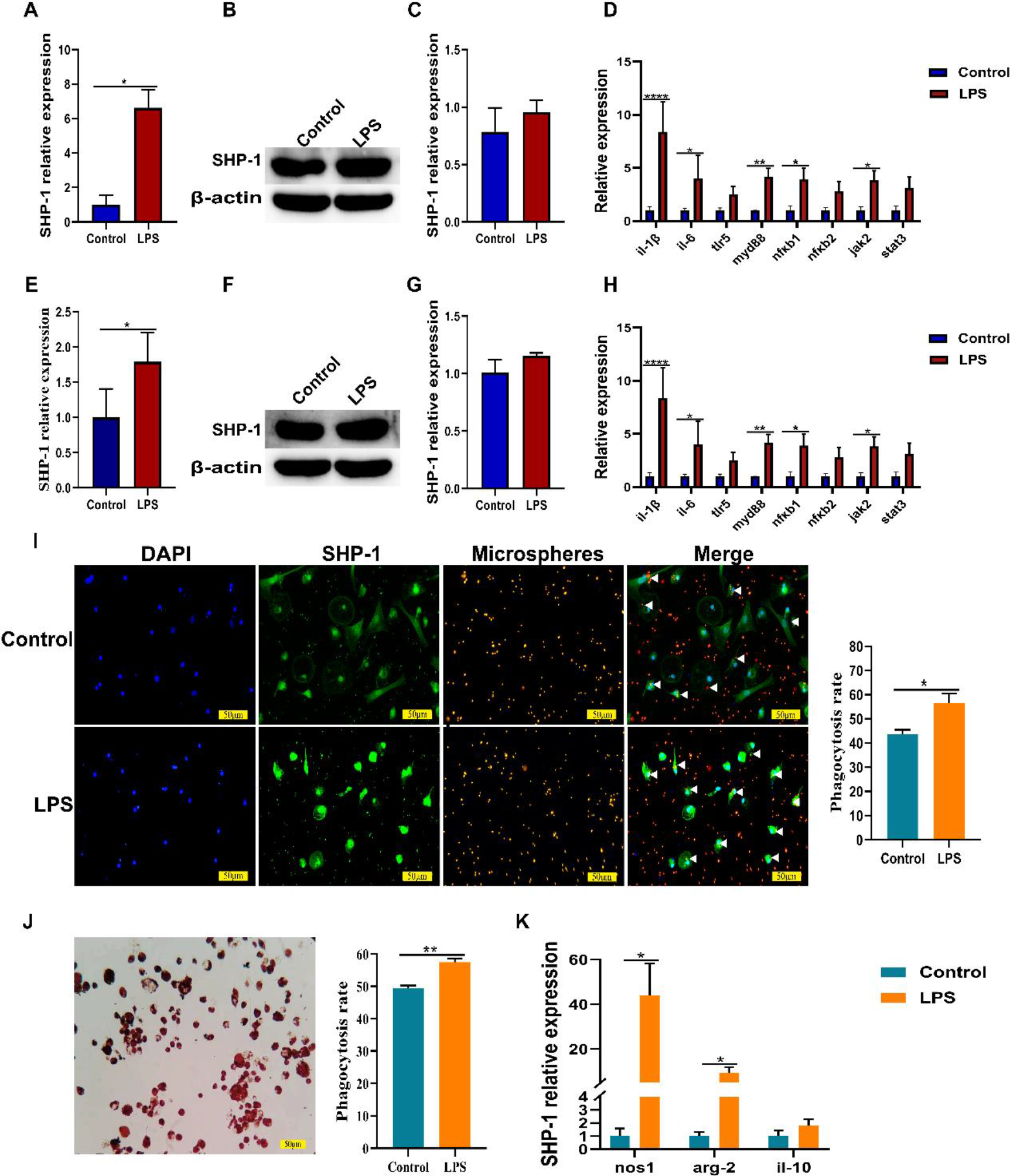
LPS induced inflammation model of hepatocytes and macrophages of Chinese tongue sole and effected macrophages polarization and phagocytic activity. (**A to C**) Expression of SHP-1 in hepatocytes after LPS stimulation. (**D**) Expression of *il-1β*, *il-6*, *tlr5*, *myd88*, *nfκb1*, *nfκb2*, *jak2*, and *stat3* in hepatocytes treated or untreated with LPS. (**E to G**) Expression of SHP-1 in macrophages after LPS stimulation. (**H**) Expression of *il-1β*, *il-6*, *tlr5*, *myd88*, *nfκb1*, *nfκb2*, *jak2*, and *stat3* in macrophages treated or untreated with LPS. (**I**) Macrophage phagocytosis determined by measuring the number of macrophages that engulfed fluorescent microspheres via immunofluorescence experiments. DAPI is used to stain the cell nucleus as blue fluorescence. SHP-1 displays green fluorescence, which is located in the nucleus and cytoplasm of macrophages, by specifically binding to the CoraLite488-conjugated goat anti-rabbit IgG secondary antibody; The fluorescent microspheres can be excited by a wavelength of 565 nm to produce red fluorescence. (**J**) Phagocytosis measured by the absorbance of neutral red engulfed by macrophages. (**K**) Macrophage polarization markers including *nos1*, *arg-2*, and *il-10* were determined by qRT-PCR. Data are presented as means ± SD, n = 3 biologically independent experiments [(A) to (K)]. * *P* < 0.05, ** *P* < 0.01, *** *P* < 0.001, **** *P* < 0.0001 by t test. Scale bars, 50 μm [(I) and (J)].

There are two major subpopulations of macrophages with distinct functions, including classically activated or inflammatory (M1) and alternatively activated or anti-inflammatory (M2) macrophages, the phenomenon of which termed “macrophage polarization” (*31–33*). Our results showed that both M1 marker genes (*il-1β*, *il-6*, and *nos1*) and M2 marker genes (*arg-2* and *il-10*) were all increased, but to different extents, indicating that primary macrophages were predominantly polarized into M1 macrophages (Fig. 3, H and K). Moreover, LPS promoted the phagocytic activity of macrophages using phagocytose fluorescent microspheres and neutral red (Fig. 3, I and J).

### SHP-1 overexpression inhibits LPS induced inflammatory response and affects macrophages polarization and phagocytosis

To verify the function of SHP-1 in inflammatory response *in vivo*, we performed *in vitro* analyses by transfecting SHP-1 expression plasmid into hepatocytes and macrophages. Results of fluorescence microscope imaging, qRT-PCR, and Western blot indicated that SHP-1 was successfully transfected into hepatocytes and macrophages, respectively (Fig. 4, A to H). With the overexpression of SHP-1, the expressions of *il-1β*, *il-6*, *tlr5*, *myd88*, *nfκb1*, *nfκb2*, *jak2*, and *stat3* were significantly suppressed (Fig.4, I and J). These results indicated that SHP-1 can significantly inhibit inflammatory responses in both hepatocytes and macrophages.

**Fig. 4.**
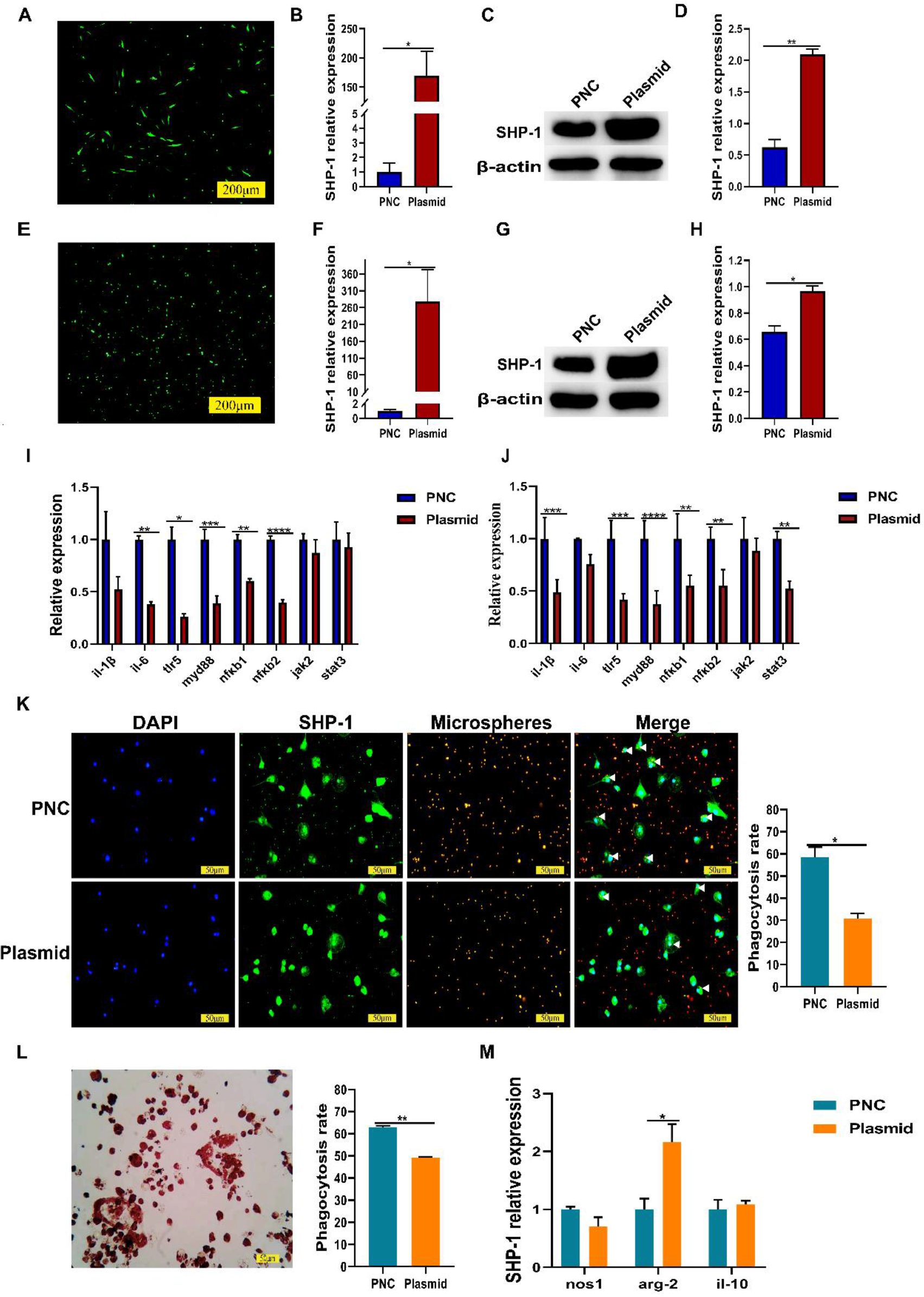
SHP-1 overexpression plasmid inhibited LPS induced inflammatory response and effected macrophages polarization and phagocytic activity. (**A to H**) Overexpression of SHP-1 in hepatocytes and macrophages validated by fluorescence microscopy imaging, qRT-PCR, and Western blot. Plasmid indicates transfection of SHP-1 plasmid; PNC indicates transfection of empty vector as negative control. **(I and J**) Expression of *il-1β*, *il-6*, *tlr5*, *myd88*, *nfκb1*, *nfκb2*, *jak2*, and *stat3* in hepatocytes and macrophages which were transfected with SHP-1 plasmid or empty vector followed by LPS stimulation. (**K**) Macrophage phagocytosis determined by measuring the number of macrophages that engulfed fluorescent microspheres via immunofluorescence experiments. DAPI is used to stain the cell nucleus as blue fluorescence. SHP-1 displays green fluorescence, which is located in the nucleus and cytoplasm of macrophages, by specifically binding to the CoraLite488-conjugated goat anti-rabbit IgG secondary antibody; The fluorescent microspheres can be excited by a wavelength of 565 nm to produce red fluorescence. (**L**) Phagocytosis measured by the absorbance of neutral red engulfed by macrophages. (**M**) Macrophage polarization markers including *nos1*, *arg-2*, and *il-10* were determined by qRT-PCR. Data are presented as means ± SD, n = 3 biologically independent experiments [(A) to (M)]. * *P* < 0.05, ** *P* < 0.01, *** *P* < 0.001, **** *P* < 0.0001 by t test. Scale bars, 50 μm [(K) and (L)].

Based on the overexpression of SHP-1 inhibiting the progress of inflammatory response, we further explored whether SHP-1 can affect the phagocytic activity and polarization of macrophages. Both macrophages phagocytosis of fluorescent microspheres and neutral red experiments demonstrated that SHP-1 overexpression suppressed the macrophage phagocytosis (Fig. 4, K and L). qRT-PCR results showed that SHP-1 overexpression inhibited the expression of *il-1β*, *il-6*, and *nos1*, but increased the expression of *arg-2* and *il-10* (Fig. 4, I, J and M). The above results indicated SHP-1 could inhibit inflammatory response through affect polarization of macrophages.

### SHP-1 siRNA and inhibitor further promote LPS induced inflammatory response and affect macrophages polarization and phagocytosis

To further verify the function of SHP-1 in inflammation, we artificially downregulated its expression *in vitro*. Firstly, we used *shp-1* siRNA to transfect hepatocytes and macrophages separately. Results of fluorescence microscope imaging, qRT-PCR, and Western blot showed that *shp-1* siRNA was successfully transfected into hepatocytes and macrophages, respectively (Fig. 5, A to H). The reduction of SHP-1 expression correlated with the increased expression of *il-1β*, *il-6*, *tlr5*, *myd88*, *nfκb1*, *nfκb2*, *jak2*, and *stat3* (Fig. 5, I and J). Macrophage phagocytosis of fluorescent microspheres and neutral red experiments demonstrated that downregulation of SHP-1 further enhanced the phagocytic activity of macrophages (Fig. 5, K and L). qRT-PCR results showed that *shp-1* siRNA increased the expression of *il-1β*, *il-6*, *nos1*, *arg-1*, and *il-10* (Fig. 5, I, J and M). Furthermore, SHP-1 inhibitor was used to verify the effects of *shp-1* siRNA. The inhibitor successfully suppressed the SHP-1 expression in hepatocytes and macrophages (Fig. 6, A to C, E to G), and it showed the same regulatory effect as siRNA (Fig. 6, D, H, I to K).

**Fig. 5.**
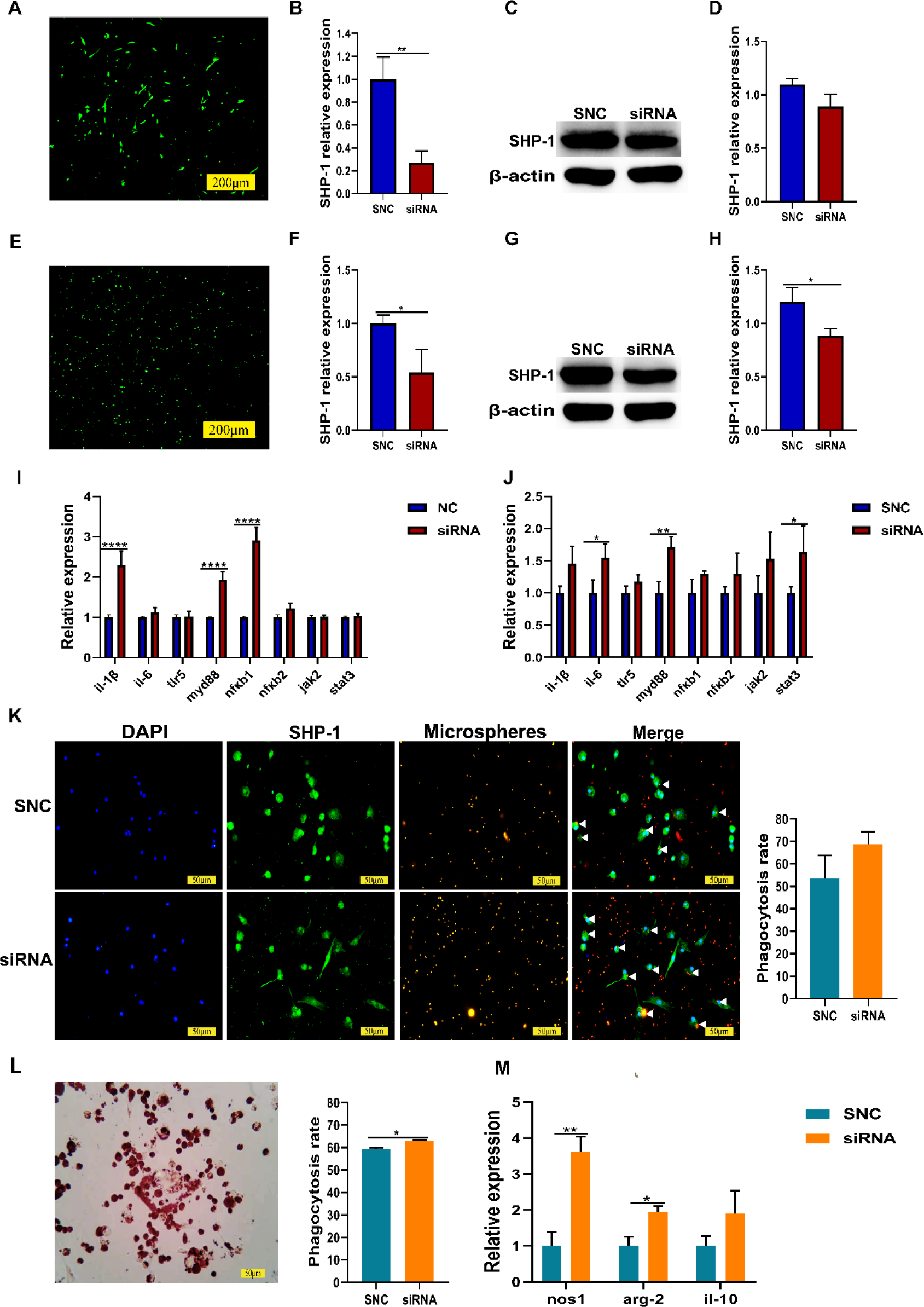
SHP-1 siRNA promoted LPS induced inflammatory response, phagocytic activity of macrophages and effected macrophages polarization. (**A to H**) Silencing SHP-1 in hepatocytes and macrophages validated by fluorescence microscopy imaging, qRT-PCR, and Western blot. siRNA indicates transfection of *shp-1* siRNA; SNC indicates transfection of randomly synthesized siRNA as negative control. (**I and J**) Expression of *il-1β*, *il-6*, *tlr5*, *myd88*, *nfκb1*, *nfκb2*, *jak2*, and *stat3* in hepatocytes and macrophages which were transfected with *shp-1* siRNA or randomly synthesized siRNA followed by LPS stimulation. **(K**) Macrophage phagocytosis determined by measuring the number of macrophages that engulfed fluorescent microspheres via immunofluorescence experiments. DAPI is used to stain the cell nucleus as blue fluorescence. SHP-1 displays green fluorescence, which is located in the nucleus and cytoplasm of macrophages, by specifically binding to the CoraLite488-conjugated goat anti-rabbit IgG secondary antibody; The fluorescent microspheres can be excited by a wavelength of 565 nm to produce red fluorescence. (**L**) Phagocytosis measured by the absorbance of neutral red engulfed by macrophages. (**M**) Macrophage polarization markers including *nos1*, *arg-2*, and *il-10* were determined by qRT-PCR. Data are presented as means ± SD, n = 3 biologically independent experiments [(A) to (M)]. * *P* < 0.05, ** *P* < 0.01, *** *P* < 0.001, **** *P* < 0.0001 by t test. Scale bars, 50 μm [(K) and (L)].

**Fig. 6.**
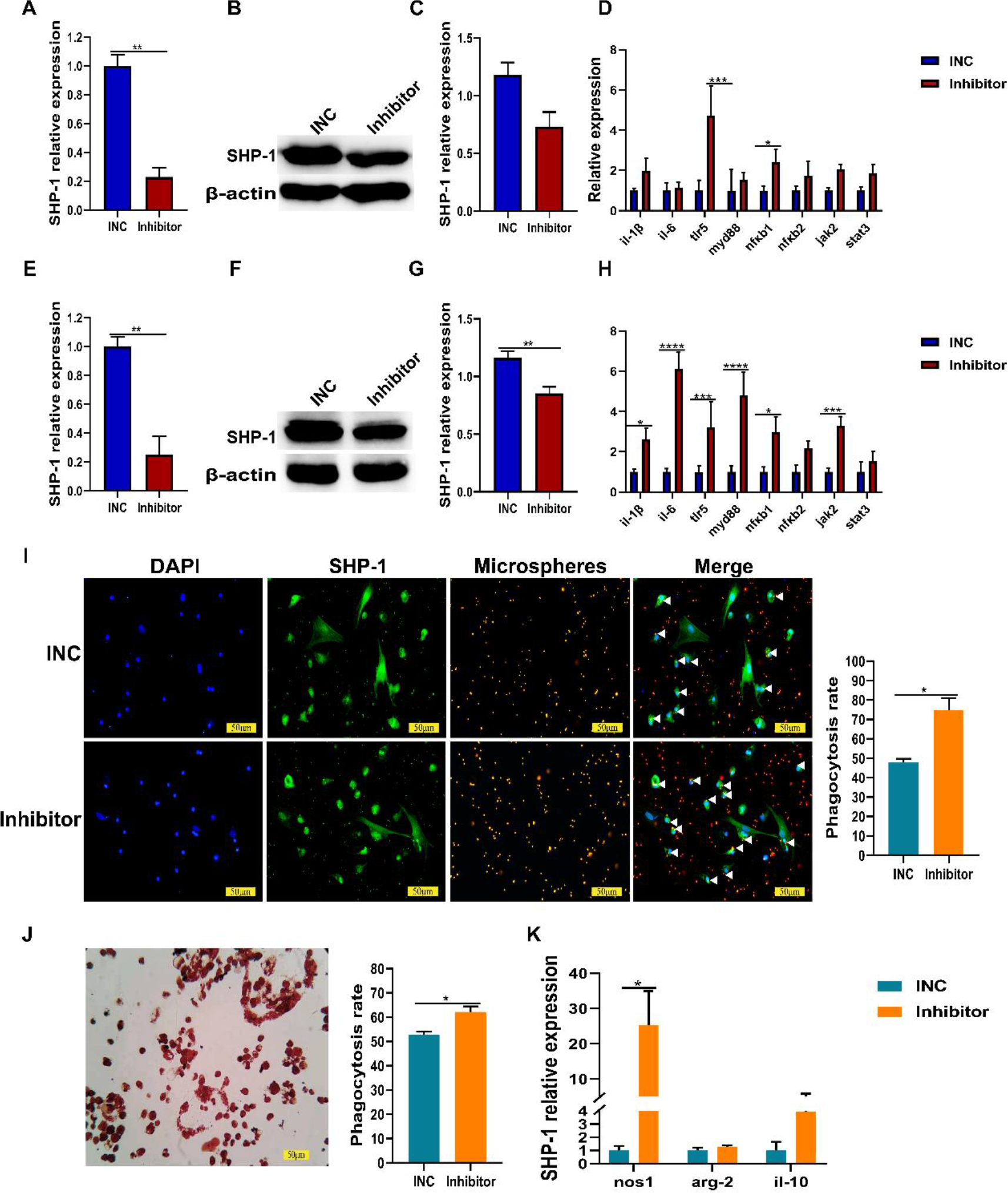
SHP-1 inhibitor promoted LPS induced inflammatory response, phagocytic activity of macrophages and effected macrophages polarization. (**A to C**) SHP-1 inhibition in hepatocytes validated by qRT-PCR and Western blot. (**D**) Expression of *il- 1β*, *myd88*, *nfκb1*, *nfκb2, il-6*, *jak2*, *stat3*, and *tlr5* in hepatocytes which were incubated with SHP-1 inhibitor or PBS followed by LPS stimulation. (**E to G**) SHP-1 inhibition in macrophages validated by qRT-PCR and Western blot. Inhibitor indicated incubation with SHP-1 inhibitor; INC indicated incubation with PBS as negative control. (**H**) Expression of *il-1β*, *myd88*, *nfκb1*, *nfκb2, il-6*, *jak2*, *stat3*, *tlr5* in macrophages which were incubated with SHP-1 inhibitor or PBS followed by LPS stimulation. (**I**) Macrophage phagocytosis determined by measuring the number of macrophages that engulfed fluorescent microspheres via immunofluorescence experiments. DAPI is used to stain the cell nucleus as blue fluorescence. SHP-1 displays green fluorescence, which is located in the nucleus and cytoplasm of macrophages, by specifically binding to the CoraLite488-conjugated goat anti-rabbit IgG secondary antibody; The fluorescent microspheres can be excited by a wavelength of 565 nm to produce red fluorescence. (J) Phagocytosis measured by the absorbance of neutral red engulfed by macrophages. (K) Macrophage polarization markers including *nos1*, *arg-2*, and *il-10* were determined by qRT-PCR. Data are presented as means ± SD, n = 3 biologically independent experiments [(A) to (K)]. * *P* < 0.05, ** *P* < 0.01, *** *P* < 0.001, **** *P* < 0.0001 by t test. Scale bars, 50 μm [(I) and (J)].

### SHP-1 protects Chinese tongue sole and its macrophages from oxidative stress

Next, we sought to determine whether SHP-1 can protect Chinese tongue sole against oxidative stress damage by evaluating the respiratory burst activity and measuring two important indicators of oxidative stress status including superoxide dismutase (SOD) activity and malondialdehyde (MDA) level. We first showed that LPS stimulated the respiratory burst activity of macrophages (Fig. 7A). SHP-1 overexpression significantly reduced the LPS induced respiratory burst activity compared with negative control (Fig. 7B), while reduction of SHP-1 using siRNA or inhibitor significantly promoted respiratory burst activity of macrophages (Fig. 7, C and D). Furthermore, LPS-stimulated macrophages and *V. anguillarum*-infected Chinese tongue sole exhibited increased MDA level but decreased SOD activity (Fig. 7, A, E and F). When SHP-1 was overexpressed in macrophages and Chinese tongue sole, MDA level was significantly decreased and SOD level was significantly increased compared with negative control (Fig. 7, B, E and F). In contrast, SHP-1 inhibition further increased MDA level and decreased SOD activity (Fig. 7, C to F). These results indicate that SHP- 1 has the capability to mitigate oxidative stress.

**Fig. 7.**
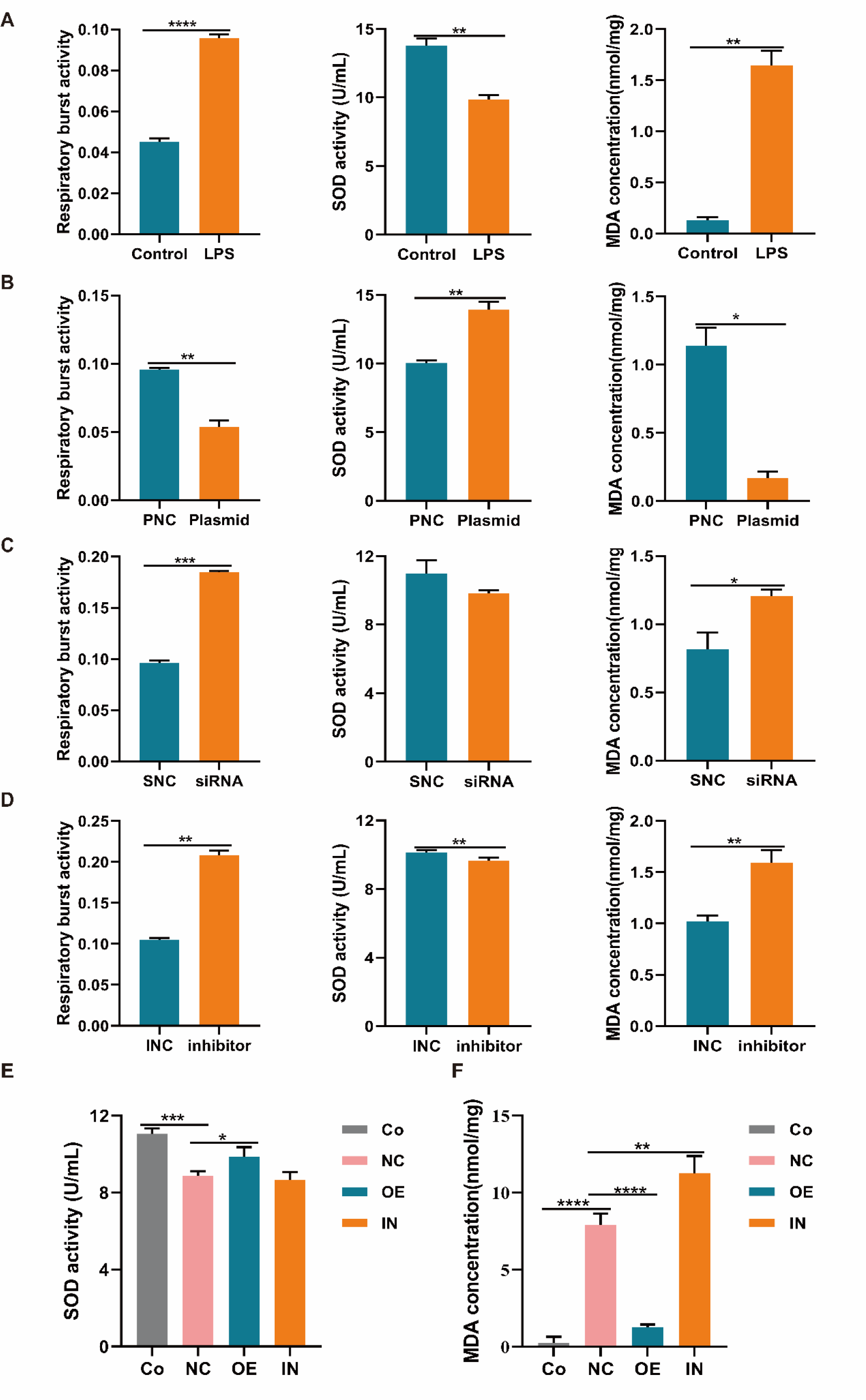
SHP-1 protected Chinese tongue sole from oxidative stress. (**A**) The respiratory burst activity, SOD and MDA in macrophages treated or untreated with LPS. (**B**) The respiratory burst activity, SOD and MDA in macrophages transfected with *shp- 1* plasmid or empty vector group. (**C**) The respiratory burst activity, SOD and MDA in macrophages transfected with *shp-1* siRNA or randomly synthesized siRNA. (**D**) The respiratory burst activity, SOD and MDA in macrophages treated with SHP-1 inhibitor or PBS. (**E to F**) The SOD and MDA level in the liver of Chinese tongue sole in different groups. CO, blank control group; NC, PBS injection followed by *V. anguillarum* infection; OE, SHP-1 overexpression followed by *V. anguillarum*; IN, SHP-1 inhibition followed by *V. anguillarum* infection. Data are presented as means ± SD, n = 3 biologically independent experiments [(A) to (F)]. [(A) to (D)] * *P* < 0.05, ** *P* < 0.01, *** *P* < 0.001, **** *P* < 0.0001 by t test. [(E) to (F)] * *P* < 0.05, ** *P* < 0.01, *** *P* < 0.001, **** *P* < 0.0001 by one-way ANOVA.

### SHP-1 inhibits the phosphorylation and nuclear translocation of NFκB1 to suppress the NFκB signaling

Upon infections, the canonical NFκB signaling pathway is activated, followed by degradation of IκB protein and release of NFκB dimer (NFκB1/Rela), which undergoes post-translational modifications such as phosphorylation, and then translocated into the nucleus to induce the transcription of target genes (*34, 35*). Due to these, we first measured the phosphorylation of NFκB1 and expression changes of immune-related molecules by Western blot. The results showed the phosphorylation of NFκB1 was markedly increased and the expression of IL-1β, IL-6, TLR5, and MYD88 were significantly induced after infection with *V. anguillarum* (Fig. 8, A and B). In addition, the overexpression and inhibition of SHP-1 respectively decreased and increased the phosphorylation of NFκB1 and the expression of selected immune-related proteins.

**Fig. 8.**
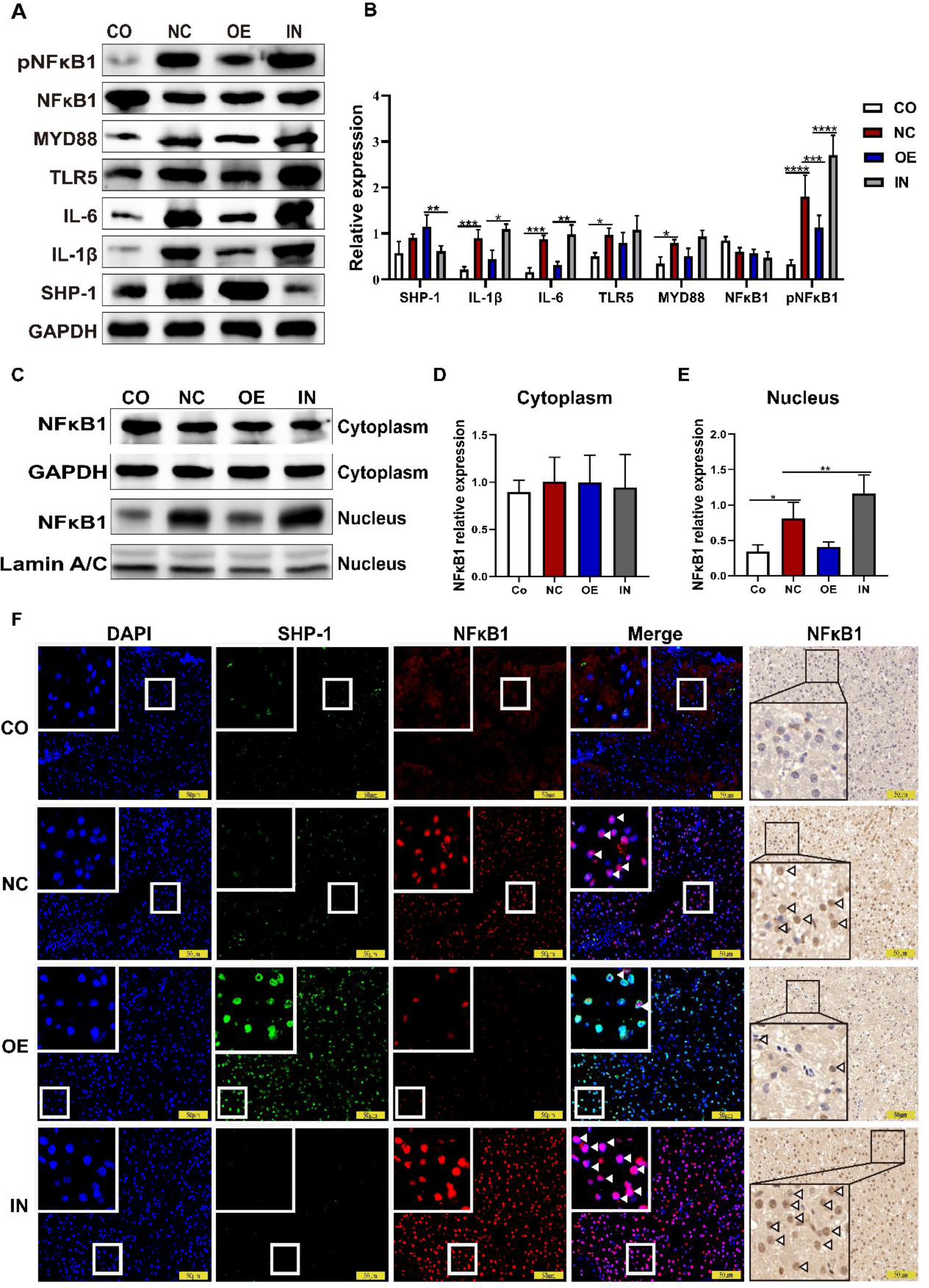
SHP-1 inhibits the phosphorylation and nuclear translocation of NFκB1 to suppress NFκB signaling pathway. (**A and B**) Expression of SHP-1, IL-1β, IL-6, TLR5, MYD88, NFκB1, pNFκB1 in the liver of Chinese tongue sole after the infection of *V. anguillarum* (LD80) by Western blot. CO, blank control group; NC, PBS injection followed by *V. anguillarum* infection; OE, SHP-1 overexpression followed by *V. anguillarum*; IN, SHP-1 inhibition followed by *V. anguillarum* infection. (**C to E**) NFκB1 protein content in cytoplasm and nucleus detected by Western blot. The internal references are GAPDH and Lamin A/C respectively. (**F**) Immunofluorescence and immunohistochemistry analysis of NFκB1 of different groups. DAPI is used to stain the cell nuclei with blue fluorescence; The SHP-1 combined with CoraLite488- conjugated goat anti-rabbit IgG secondary antibody shows green fluorescence; NFκB1 combined with Cy5 goat anti-rabbit IgG secondary antibody displays red fluorescence; Brown represents a positive immunohistochemical result and cell nucleus was stained blue. White arrow represents NFκB1 in the nucleus. Data are presented as means ± SD, n = 3 biologically independent experiments [(A) to (E)]. * *P* < 0.05, ** *P* < 0.01, *** *P* < 0.001, **** *P* < 0.0001 by one-way ANOVA. Scale bars, 50 μm (F).

Meanwhile, we investigated whether SHP-1 can affect the nuclear translocation of NFκB1. To this end, proteins in the cytoplasm and nucleus were separately isolated to examine the distribution of NFκB1. No significant difference was observed among different groups in cytoplasm, whereas the NFκB1 protein was differentially expressed between groups in the nucleus. Compare with the negative control, SHP-1 overexpression reduced NFκB1 content, while SHP-1 inhibition significantly increased its presence (Fig. 8, C to E). To further elucidate the regulatory mechanism of SHP-1 acting on NFκB signaling pathway, we performed immunofluorescence co-localization and immunohistochemical analysis. Microscopic imaging showed similar results to Western blotting (Fig. 8F), further proving that SHP-1 inhibits the nuclear translocation of NFκB1. Overall, we demonstrate for the first time that SHP-1 not only inhibits NFκB1 phosphorylation but also nuclear translocation.

### SHP-1 regulates the NFκB signaling pathway by interacting with NFκB1

In view of the effect of SHP-1 on NFκB signaling pathway, we speculated that there was an interactive relationship between SHP-1 and NFκB1. To test the hypothesis, we first examined the interaction of SHP-1 with NFκB1 through Co-IP. Results showed that SHP-1 can indeed interact with NFκB1 (Fig. 9A). Furthermore, we detected the localization of SHP-1 and NFκB1 in hepatocytes through immunofluorescence co- localization experiments. The results showed that SHP-1 and NFκB1 were expressed in both the nucleus and cytoplasm, and there was obvious co-localization between the two in the cytoplasm (Fig. 9B), which further support the interaction between SHP-1 and NFκB1.

**Fig. 9.**
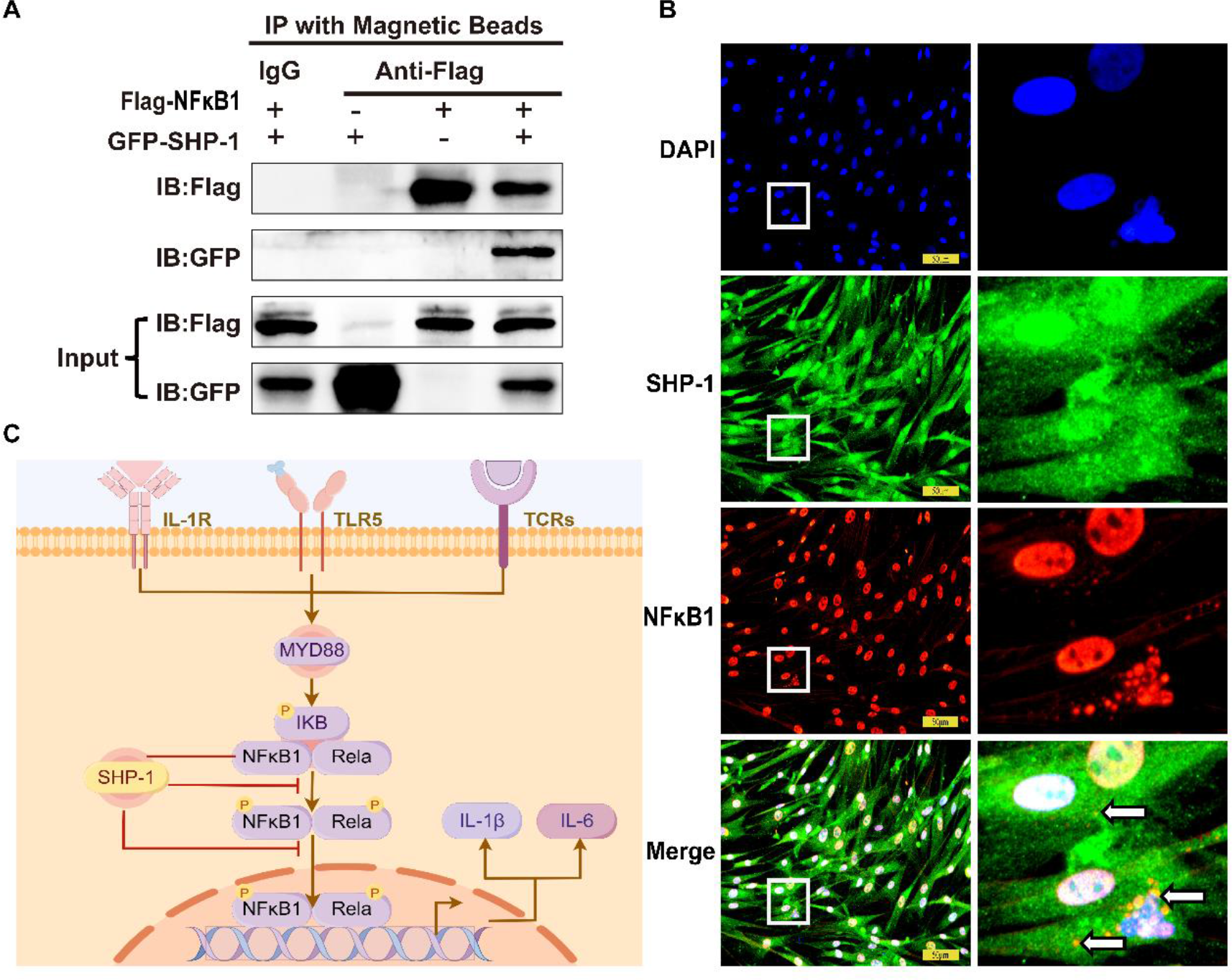
SHP-1 inhibits the NFκB signaling pathway by interacting with NFκB1. (**A**) Co-immunoprecipitation experiment used to examine the interaction between SHP-1 and NFκB1. Co-IP was performed with Flag-labeled magnetic beads. (**B**) Immunofluorescence co-localization analysis of SHP-1 and NFκB1 in hepatocytes, blue fluorescence represents nucleus, green fluorescence represents SHP-1 proteins, red fluorescence represents NFκB1 proteins, Merge represents combining three fluorescent images together. DAPI is used to stain the cell nucleus as blue fluorescence; SHP-1 combined with CoraLite488-conjugated goat anti-rabbit IgG secondary antibody shows green fluorescence; NFκB1 combined with Cy5 goat anti-rabbit IgG secondary antibody shows red fluorescence; White arrow represents SHP-1 and NFκB1 co- localization. (**C**) Schematic diagram of SHP-1 regulating NFκB signaling pathway. Mechanisms revealed in this study are highlighted in red. Scale bars, 50 μm (B).

## Discussion

SHP-1 has emerged as a key regulator in a myriad of signaling cascades, affecting various biological processes. Specifically, it plays a pivotal part in inflammation and immune response, however, the underlying mechanism is not fully understood. In this study, inspired by our previous study (*28*) and literature reading, we determined to reveal the function and underlying mechanism of SHP-1 on inflammation caused by bacterial infection, using both teleost model and cellular model of Chinese tongue sole. We demonstrated that SHP-1 exert negative roles to avoid excessive inflammation and immune response to protect the body from oxidative stress, tissue damage, and even death, manifested by its molecular functions on the expression of inflammatory cytokines, key immune-related genes, and physiological indicators. The highlight of this study is that we discovered a novel mechanism by which SHP-1 negatively regulate the NFκB signaling, i.e., SHP-1 inhibits NFκB1 phosphorylation and nuclear translocation by interacting with NFκB1 (Fig. 9C). Our findings shed light on the critical role of SHP-1 in regulating inflammatory responses and open new avenues for therapeutic intervention for the treatment of inflammation-related diseases, including cancer and infectious diseases.

The present finding that SHP-1 reduced mortality and histopathological deterioration following *V. anguillarum* infection suggests a protective role for SHP-1 in the immune response irrespective of the bacterial concentration (LD80 or LD50). The results were intriguing, which may explain why SHP-1 showed differential expression after the same bacterial infection in the same teleost in our previous study (*36*), i.e., SHP-1 underwent dynamic regulation to maintain the cell and tissue homeostasis to avoid death after infection. These results are consistent with its effects in mammals, whereby SHP-1 attenuates corneal damage induced by LPS challenge (*37*) as well as reduces lymphocyte infiltration and epithelia hyperplasia/metaplasia in mouse stomach (*38*).

Going beyond the phenotypic observations, we investigated the molecular mechanisms underlying SHP-1’s immunomodulatory effects. After pathogen infections, immune-related signaling pathways including TLR5-MYD88-NFκB and JAK-STAT3 were activated, leading to the production and release of pro-inflammatory cytokines, including IL-1β and IL-6 (*12, 13, 16, 18*). The exaggerated activation of these signaling cascades and excessive release of inflammatory cytokines is the central mechanism that initiates cytokine storm, damage to tissues and organs, and even death (*15, 21*). For instance, studies of coronavirus disease 2019 (COVID-19) showed that uncontrolled activation of NFκB and JAK-STAT3 signaling in immune cells upon the stimulation of severe acute respiratory syndrome coronavirus (SARS-CoV-2) is responsible for the hyper-production of inflammatory mediators and severe acute respiratory distress syndrome (ARDS) in human (*39*). In our present study, both *in vivo* fish model and *in vitro* cellular model demonstrated that SHP-1 significantly inhibited these signaling cascades, suppressed the excessive secretion of pro-inflammatory cytokines, and increased anti-inflammatory cytokine expression, providing a preliminary insight into the regulatory mechanism of SHP-1. It is worth noting that NFκB signaling was the most significantly changed among the studied pathways.

NFκB is a well-known family of transcription factors, serving as a molecular lynchpin to infection, inflammation, and cancer (*35*). NFκB proteins normally exist in the cytoplasm and bind to the inhibitory protein IκB, thereby being maintained in an inactivated state. Upon stimulation such as infections and cytokines, canonical signaling pathway is primarily activated, where IκB is phosphorylated and degraded, thereby releasing NFκB1/Rela dimer which subsequently translocate to the nucleus and activate target gene transcription (*34, 35*). SHP-1 regulates NFκB signaling pathway through Rela has been studied (*9*), where SHP-1 suppressed the proliferation and metastasis of hepatoma cell via Rela dephosphorylation. Currently, its role on NFκB1 dephosphorylation and nuclear translation has been lacking. Through combined approaches (Co-IP, IF, and IHC), we demonstrated that SHP-1 could inhibit the phosphorylation and nuclear translocation of NFκB1 by interacting with NFκB1. To our limited knowledge, this is the first study showing the regulatory mechanism of SHP-1 on NFκB1. We believe this is a significant discovery since our results link the phenotype, cellular function, and intrinsic mechanism.

Macrophages, as the main effector cells of innate immunity, play an important role in the body’s immune defense, immune homeostasis, and immune surveillance (*40, 41*). They can undergo polarization and functional changes to deal with different micro- environments (*31–33*). M1/M2 macrophages balance governs the fate of an organ in inflammation (*42*). Compared with those in mammals, macrophages of teleost possess specific functions to regulate inflammatory response (*43*). Therefore, studies on teleost macrophages may help improve our understanding of its roles in inflammation. Previous studies have shown that macrophages derived from the head kidney of common carp (*Cyprinus carpio*) exhibited functional polarization under LPS and cyclic adenosine monophosphate (cAMP) stimulation (*44, 45*). In this study of Chinese tongue sole, we proved that SHP-1 can inhibit macrophages polarize into M1 phenotype and suppress the secretion of pro-inflammatory cytokines such as IL-1β and IL-6 and the activation of NFκB signaling pathway. Meanwhile, SHP-1 was shown to induce macrophages polarized into M2 phenotype and promote the secretion of anti-inflammatory cytokines such as *il-10* and *arg-2*. The results of macrophage polarization further support that SHP-1 acts as a negative regulator in inflammatory response. We will verify our results of macrophage polarization by flow cytometry in the future research.

Oxidative stress indicates an imbalance between the over-production of reactive oxygen species (ROS) and their protective elimination, which is often associated with inflammation and can contribute to the pathogenesis of various diseases (*46*). Studies have illustrated that the generation of ROS was concomitant with the activation of tyrosine phosphorylation, and some emphasized that protein tyrosine phosphatase (PTPs) may serve as oxidation sensors in the context of phosphotyrosine signaling (*47*). In the present study, we explored whether SHP-1 could affect oxidative stress in Chinese tongue sole by examining the macrophages respiratory burst activity and oxidative stress indicators (SOD and MDA). Our finding showed that SHP-1 significantly inhibited the oxidative stress of Chinese tongue sole both *in vivo* and *in vitro*, consistent with previous studies (*48*, *49*). The observation that SHP-1 protects against oxidative stress adds another dimension to its role in maintaining cellular and individual homeostasis.

Collectively, the findings presented in this study provide valuable insights into the versatile functions and mechanisms of SHP-1 in the negative regulation of inflammation/immune response. The identification of SHP-1 as a modulator of NFκB signaling and the newly revealed mechanisms offers therapeutic opinions for the treatment of cancer and bacterial diseases. Future studies are necessary to elucidate the mechanisms more comprehensively and fully explore the potential of SHP-1-based therapeutics for inflammation-associated diseases.

## Materials and Methods

### Experimental fish and experimental design

Chinese tongue sole (weight 10 ± 5 g, length 10 ± 3 cm) was purchased from Laizhou Mingbo Aquatic Co., Ltd. All fish were acclimated in aerated seawater (temperature 24°C, salinity 30‰, dissolved oxygen content 6.1-6.7 mg/L, pH 7.8-8.2) in a 50 L tank prior to the experiment, as previously described (*50*). The fish were divided into CO group without any treatment, NC group where PBS was injected intravenously followed by *V. anguillarum* infection, OE group where 5 μg/g SHP-1 overexpression plasmid was injected intravenously followed by *V. anguillarum* infection, and IN group where 3 μg/g SHP-1 inhibitor was injected intravenously followed by *V. anguillarum* infection. After 48 h, except the CO group, other groups were intraperitoneally separately injected with 3.7 × 10^8^ CFU/mL (LD80) and 1.6 × 10^8^ CFU/mL (LD50) of *V. anguillarum*. The *V. anguillarum* used in the experiment was verified by 16S rDNA sequencing. The bacterium was incubated to mid-logarithmic stage at 28°C in 2216E medium, and the concentration was determined by the colony-forming unit (CFU) method. Samples of liver, intestine, spleen, and gill were collected at each time point following the infection (0 h, 12 h, 24 h, 48 h, and 72 h), snap frozen in liquid nitrogen, and stored at -80°C for subsequent RNA and protein extraction. The fish were anesthetized with MS-222 before injection and sampling, and all the animal experiments were performed in accordance with the guidelines approved by the Ethics Committee of Qingdao University.

### Cell culture

Hepatocyte cell line of Chinese tongue sole was from China Yellow Sea Fisheries Research Institute, cultured with 15% fetal bovine serum (FBS), 0.0028% mercaptoethanol, 0.2% epidermal growth factor (EGF), 0.2% basic fibroblast growth factor (bFGF), 0.1% leukemia inhibitory factor (LIF), 1% triple antibiotics (penicillin, streptomycin, and amphotericin B), and L15 culture medium. Cells were cultured in an atmosphere at 24°C.

Extraction and purification of primary macrophages were modified and optimized based on a previous protocol (*51, 52*), and its effectiveness was verified by our group.

In brief, 30 mL 3% soluble starch dissolved in LB broth was sterilized at 120°C and injected intraperitoneally to elicit the macrophages. One day later, the peritoneal cavity of the fish was lavaged with 50 mL of L15 medium to obtain the macrophages. The culture conditions of primary macrophages were consistent with those of hepatocytes. The human embryonic kidney cell line (HEK293T) was obtained from ATCC (Rockville, USA), cultured with 10% FBS, 1% triple antibody and 89% DMEM culture medium. Cells were cultured in an atmosphere of 5% CO2 at 37°C.

### qRT-PCR assay

The expression of *shp-1* in tissues and cells were determined by quantitative real time polymerase chain reaction (qRT-PCR). Total RNA was extracted by spark easy cell RNA kit (Solarbio, China) and reverse transcribed into cDNA by HiScript III RT Super Mix (Vazyme, China). The primers are shown in Table 1, using *18S rRNA* gene as the internal reference. qRT-PCR was performed on the ABI 7500 fast quantitative real-time amplification instrument (ABI, USA), and the reaction procedure was as follows: 95°C for 30 s; 95°C for 10 s, 60°C for 30 s, 40 cycles. Three biological replicates were set for each sample, and the 2^-ΔΔCt^ method was used to calculate the relative expression.

**Table 1.**
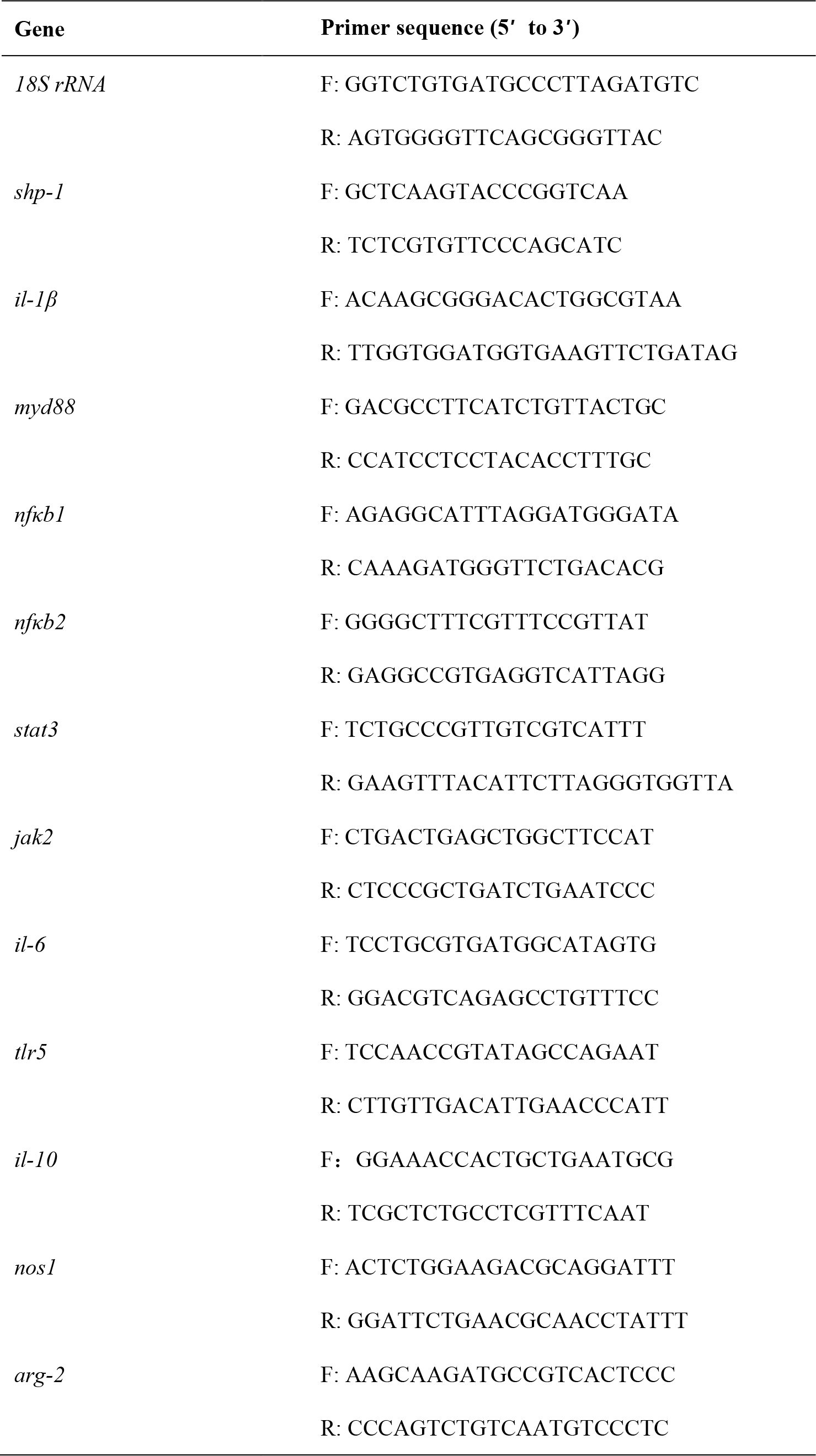
Primers used in the study for qRT-PCR.

### Western blot assay

At 12 h post the infection of *V. anguillarum*, liver tissues of different experimental groups were collected. The total protein was extracted by RAPI lysate (Beyotime, China), separated by sodium dodecyl sulfate polyacrylamide gel electrophoresis (SDS-PAGE), and then transferred on polyvinylidene fluoride (PVDF) membrane. The membrane was probed with rabbit anti-SHP-1 polyclonal antibody, rabbit anti-IL-6 polyclonal antibody, rabbit anti-IL-1β polyclonal antibody (ABclonal, China), rabbit anti-MYD88 polyclonal antibody, mouse anti-NFκB1 monoclonal antibody (Proteintech, China), phospho-NFκB1-S337 polyclonal antibody (ABclonal, China). Meanwhile, the same amount of extract was recognized using recombinant anti-GAPDH antibody (Servicebio, China) as the comparative control. HRP-conjugated goat anti-rabbit IgG and HRP-conjugated goat anti-mouse IgG were used as secondary antibody (ABclonal, China). The blots were visualized using enhanced chemiluminescence (ECL) (Beyotime, China) and imaged on a luminescence imaging system (Bio-Rad, USA). The SHP-1, IL-6, TLR5, MYD88 polyclonal antibodies of Chinese tongue sole were synthesized by GenScript (China).

### Hematoxylin-eosin (HE) staining

The liver and intestine tissues of Chinese tongue sole at different time points post infection were collected, and HE staining was performed to detect the histopathological changes. The tissues were cut into 0.5 × 0.5 cm tissue blocks and fixed with 4% paraformaldehyde. The fixed tissue samples were embedded in paraffin, and they were cut into sections with a thickness of about 4-8 μm using a rotary microtome. The prepared films were dewaxed, then stained with hematoxylin and eosin respectively, and finally the stained films were blocked with neutral resin.

### Immunofluorescence assay

Hepatocytes and primary macrophages from Chinese tongue sole were washed with pH 7.4 phosphate buffered saline (PBS) and fixed with 4% paraformaldehyde, followed by permeabilization with 0.5% Triton X-100. The cell plates were then incubated with rabbit anti-SHP-1 IgG (20 μg/mL), mouse anti-NFκB1 IgG (20 μg/mL), or normal rabbit serum (1:100), followed by CoraLite488-conjugated goat anti-rabbit IgG and CoraLite564-conjugated goat anti-mouse IgG secondary antibody (Abcam, USA). Hepatocytes and primary macrophages were counterstained for nuclei with 1 μg/mL of 4’,6-Diamidino-2-phenylindole. Images were captured with fluorescence microscope (Nikon, Japan).

Paraffin sections of liver tissues were dewaxed and washed, and antigen retrieval was performed with EDTA antigen retrieval buffer. The sections were placed in 3% hydrogen peroxide solution and incubated in the dark at room temperature for 25 min to block the endogenous peroxidase, and then blocked with negative serum. For immunofluorescence double labeling experiment, paraffin sections were incubated with rabbit anti-SHP-1 IgG (20 μg/mL), mouse anti-NFκB1 IgG (20 μg/mL) or normal rabbit serum (1:100). Tyramide signal amplification (TSA) was introduced for immunofluorescence triple labeling experiment. Paraffin sections were incubated with rabbit anti-SHP-1 IgG (20 μg/mL), rabbit anti-IL-1β IgG (20 μg/mL), rabbit anti-IL-6 IgG (20 μg/mL) or normal rabbit serum (1:100), and then used CoraLite488-conjugated goat anti-rabbit IgG, Cy3-conjugated goat anti-rabbit IgG (H+L) and Cy5-conjugated goat anti-rabbit IgG (H+L) secondary antibody (1 μg/mL, Servicebio, China). Sections were counterstained for nuclei with 1 μg/mL of 4’,6-Diamidino-2-phenylindole. Scanning and imaging were accomplished with a fluorescence microscope (Nikon, Japan).

### Construction of eukaryotic expression plasmid

The coding sequence (CDS) of *shp-1* and *nfκb1* were obtained from the NCBI database (https://www.ncbi.nlm.nih.gov/). Primers of *shp-1* were designed (Forward: CGGCTAGCATGGTTCGGTGGTTCCAC, Reverse: CCAAGCTTTTTTTTTCGCAACGAGCCGC). The cDNA of *shp-1* was obtained by reverse transcription of liver total RNA and subcloned into Hind III/BamH I (Takara, Japan) sites of pcDNA3.1EGFP vector. The DNA encoding *nfκb1* synthesized by Shanghai Genomics Institute was subcloned into Hind III/Xho I (Takara, Japan) sites of pcDNA3.1-3xFlag- C vector, respectively. Plasmids was extracted using an endotoxin-free plasmid extraction kit (TIANGEN, China).

### Silence of SHP-1 expression

The siRNA targeting *shp-1* was designed and performed by Research Cloud (Jinan, China). The siRNA duplex with sense sequence (5’-GAAACCUGAAGCUCAAACA-3’) and anti-sense sequence (5′-CUUUGGACUUCGAGUUUGU-3’) were synthesized and conjugated with FITC. The siRNA was added to 1 × 10^6^ hepatocytes and primary macrophages to reach 100 nM in a total volume of 2 mL L15 medium and incubated for 24 h. The random-synthesized siRNA duplex with sense sequence (5’- UUCUCCGAACGUGUCACGUTT-3’) and anti-sense sequence (5’-ACGUG ACACGUUCGGAGAATT-3’) was used for control.

The SHP-1 inhibitor named TPI (Topscience, China) was added to hepatocytes and primary macrophages to reach 5 μmol in a total volume of 2 mL L15 medium and incubated for 24 h.

### Transfection

The recombinant plasmid or siRNA was added into a sterile centrifuge tube, then the equal dose of zeta transfection reagent (Life Technologies, USA) was added and mixed, followed by co-incubation for 15 min. The premix was added into hepatocytes and primary macrophages, culturing for 24 h.

### Assay of phagocytic activity of primary macrophages

Phagocytosis of fluorescent microspheres or neutral red was utilized to evaluate the phagocytic activity of macrophages. At 24 h post LPS stimulation, fluorescent microspheres were added to primary macrophages in different experimental groups and incubated for 2 h, and then immunofluorescence detection was performed. Images were captured with a fluorescence microscope (Nikon, Japan). The macrophage phagocytosis rate is equal to the number of macrophages engulfing fluorescent microspheres divided by the total number of macrophages. In neutral red test, 0.01% neutral red solution (Beyotime, China) was added into macrophages of different experimental groups and incubated for 2 h. Macrophages were washed three times with PBS, lysis buffer was added for 30 min, and the absorbance value at 562 nm was read with a microplate reader (TECAN, Switzerland). The phagocytic index was represented by the absorbance value in the experimental wells divided by the absorbance value of the control wells.

### Assay of respiratory burst of primary macrophages

Nitroblue tetrazolium (NBT) assay was used to determine the production of superoxide anion for the measurement of respiratory burst activity in macrophages. In brief, 100 μL of macrophages (5 × 10^4^ cells/mL) were placed in 96-well plate for 24 h. SHP-1 overexpression plasmid, siRNA, inhibitor, and negative control were transfected into macrophages respectively and incubated for 48 h. At 24 h post LPS stimulation, NBT (Beyotime, China) was added into 96-well cell plate and incubated at 24°C for 1 h, and the supernatant was discarded. Then, 120 μL of methanol was added to terminate the reaction. The NBT deposited inside the cells were then dissolved, first by adding 120 μL of 2 M KOH and then by adding 140 μL of DMSO (Solarbio, China). The dissolved NBT solution was transferred to a 96-well plate and absorbance was read on a microplate reader (TECAN, Switzerland) at 630 nm. Respiratory burst activity in primary macrophage was quantified as NBT reduction.

### Evaluation of oxidative stress of macrophages and liver of Chinese tongue sole

SOD content was measured by a superoxide dismutase (SOD) assay kit (Solarbio, China). Proteins of macrophages and liver tissue were extracted using RIPA cracking solution (Beyotime, China). According to the instruction, the sample to be tested, enzyme working solution, and substrate application solution were successively added into the 96-well plate and were mixed well. SOD was measured at 450 nm via a microplate reader (TECAN, Switzerland).

MDA content was determined using a malondialdehyde (MDA) test kit (Solarbio, China). Proteins of macrophages and liver tissue were extracted by grinding method. According to the instructions, the standard, sample to be tested, and reagents were added into a 1.5 mL centrifuge tube, and they were mixed well. They were then kept in a water bath above 95℃ for 40 min, taken out, and cooled in running water. After that, they were centrifuged at 4000 rpm for 10 min. The reaction solution from each tube were accurately added into a new 96-well plate., and MDA was measured at 530 nm by a microplate reader (TECAN, Switzerland).

### Immunohistochemical (IHC) staining

The liver tissue was cut into 0.5 × 0.5 cm tissue blocks and fixed with 4% paraformaldehyde for 24 h. Subsequently, sections were deparaffinized, rehydrated, and subjected to antigen retrieval. After treatment with 3% H2O2 in PBS for 30 min at room temperature, the sections were incubated with the primary antibody against NFκB1 (Proteintech, China) overnight at 4°C, followed by incubation with HRP-conjugated secondary antibody (Servicebio, China). The sections were visualized using diaminobenzidine, with haematoxylin as a counterstain. All images were captured using an optical microscopy (Nikon, Japan).

### Co-immunoprecipitation assay

Cells were grown in 6-well plates (Corning, USA) and transfected with various Flag (pcDNA3.-FLAG as control) and GFP-tagged plasmids. After 48 h of culture, Flag, and GFP-tagged proteins were expressed. Cells were gently rinsed with PBS and lysed using cell lysis buffer (Beyotime, China). 100 μL of the cell lysate was isolated as an input sample for determining protein expression, and the remaining lysate was spiked with appropriate amounts of anti-Flag magnetic beads (Proteintech, USA). The lysate- magnetic bead mixture was incubated at 4°C for 24 h with gentle shaking. After incubation, the magnetic beads were collected, and rinsed with cell lysis buffer at least thrice. Proteins which interacted with Flag-tagged proteins were also precipitated. A total of 20 μL of input samples and Co-IP samples were incubated with SDS-PAGE loading buffer (Beyotime, China) for 5 min at 100°C. Western blotting was performed using anti-Flag and anti-GFP (Proteintech, USA) antibodies to determine the precipitation of the proteins of interest.

### Statistical analysis

All experiments were performed in three biological replicates. Experimental data were analyzed using GraphPad Prism8 software. Results were expressed as mean ±standard deviation. Statistical analysis was performed using t test or one-way ANOVA to determine the significance of difference between groups (* *P* < 0.05, ** *P* < 0.01, *** *P* < 0.001, **** *P* < 0.0001).

## Acknowledgments

### Funding

National Key R&D Program of China (Grant No. 2022YFD2400401)

Shandong Key R&D Program for Academician team in Shandong (Grant No. 2023ZLYS02)

Shandong Key R&D Program (Grant No. 2021LZGC028) Taishan Scholar Climbing Project of Shandong Province, China

The Innovative Team Project of Chinese Academy of Fishery Sciences (2020TD20)

## Authorship contribution

Investigation: N. W., S. T., Z. S.

Experimental operation: N. W., M. W., H. L, S. H., Z. W., J. M. Data analysis: N. W., S. T., Z. S.

Visualization: N. W., S. T., Z. S.

Writing — original draft: N. W., S. T., Z. S.

Writing — review & editing: N. W., S. T., Z. S., S. C. Funding acquisition: Z. S., S. C.

Supervision: Z. S., S. C.

## Competing Interest

No potential conflict of interest was reported by the authors.

## Notes

### Competing Interest Statement

The authors have declared no competing interest.

